# p90RSK2, a new MLCK, rescues contractility in myosin light chain kinase null smooth muscle

**DOI:** 10.1101/2023.05.22.541840

**Authors:** Jaspreet Kalra, Mykhaylo Artamonov, Hua Wang, Aaron Franke, Zaneta Markowska, Li Jin, Zygmunt S. Derewenda, Ramon Ayon, Avril Somlyo

## Abstract

**Background:** Phosphorylation of smooth muscle (SM) myosin regulatory light chain (RLC_20_) is a critical switch leading to contraction or cell migration. The canonical view held that the only kinase catalyzing this reaction is the short isoform of myosin light chain kinase (MLCK1). Auxiliary kinases may be involved and play a vital role in blood pressure homeostasis. We have previously reported that p90 ribosomal S6 kinase (RSK2) functions as such a kinase, in parallel with the classical MLCK1, contributing ∼25% of the maximal myogenic force in resistance arteries and regulating blood pressure. Here, we take advantage of a MLCK1 null mouse to further test our hypothesis that RSK2 can function as an MLCK, playing a significant physiological role in SM contractility.

**Methods:** Fetal (E14.5-18.5) SM tissues were used as embryos die at birth. We investigated the necessity of MLCK for contractility, cell migration and fetal development and determined the ability of RSK2 kinase to compensate for the lack of MLCK and characterized it’s signaling pathway in SM.

**Results:** Agonists induced contraction and RLC_20_ phosphorylation in *mylk1*^-/-^ SM, that was inhibited by RSK2 inhibitors. Embryos developed and cells migrated in the absence of MLCK. The pCa-tension relationships in WT vs *mylk1*^-/-^ muscles demonstrated a Ca^2+^-dependency due to the Ca^2+^-dependent tyrosine kinase Pyk2, known to activate PDK1 that phosphorylates and fully activates RSK2. The magnitude of contractile responses was similar upon addition of GTPγS to activate the RhoA/ROCK pathway. The Ca^2+^-independent component was through activation of Erk1/2/PDK1/RSK2 leading to direct phosphorylation of RLC_20_, to increase contraction. RSK2, PDK1, Erk1/2 and MLCK formed a signaling complex on the actin filament, optimally positioning them for interaction with adjacent myosin heads.

**Conclusions:** RSK2 signaling constitutes a new third signaling pathway, in addition to the established Ca^2+^/CAM/MLCK and RhoA/ROCK pathways to regulate SM contractility and cell migration.

## Introduction

In vascular smooth muscle (VSM), contraction is initiated by phosphorylation of the regulatory light chain (RLC_20_) of SM myosin II on Ser19 by the SM myosin light chain kinase (smMLCK, referred to hereafter as MLCK). MLCK is activated by Ca^2+^-bound calmodulin (CaM).^1,2^ In this way, Ca^2+^ influx promotes phosphorylation of RLC_20_, activating SM myosin II and its binding to actin filaments, thereby initiating cyclic ATP-dependent cross-bridge cycling. The activity of MLCK is counterbalanced by that of the RLC_20_ phosphatase (MLCP) which induces relaxation. MLCP itself is regulated by phosphorylation, specifically by the RhoA-dependent Ser/Thr kinase ROCK. The latter is activated by the small cytosolic GTPase RhoA, downstream of G-protein coupled receptors. This pathway sustains contraction as intracellular [Ca^2+^] decreases, a phenomenon known as Ca^2+^-sensitization.^3^

The widely expressed *mylk1* gene contains alternative promotors giving rise to three proteins: the long—∼220 kDa—non-muscle myosin II light chain kinase, expressed in various cell types; the short (∼130 kDa) isoform unique to SM acting on the myosin II isoform found here; and the non-catalytic kinase related protein also known as telokin (17 kDa).^4,5^ In the classical view of SM contraction, MLCK was the sole protein kinase regulating RLC_20._ Over the past few decades, strong evidence emerged that this scheme is oversimplified. For example, direct evaluation of MLCK activity in bladder SM from a CaM-sensor-MLCK transgenic mouse model demonstrated that only about 50 % of carbachol-induced force and RLC_20_ phosphorylation was accounted for by MLCK activity.^6^ Thus, additional mechanisms exist for increasing both force and RLC_20_ phosphorylation. The possibility of other auxiliary kinases regulating SM contractility is of enormous significance, because reactivity and tone of vascular SM affects vascular resistance, and consequently blood pressure (BP).^7-9^ Studies in mice with tamoxifen-induced, SM-specific partial MLCK knockout have shown that BP is significantly reduced and salt-induced hypertension is abolished.^10,11^ However, therapeutic inhibition of MLCK is not a practical option, in part because the ubiquitously expressed non-muscle myosin shares the same catalytic domain and because of its major role in contractile activity of all SM-containing hollow organs. As a result, there is ongoing research to identify auxiliary, regulatory protein kinases in VSM as potential targets for antihypertensive therapy, as even small reductions of BP can have clinically significant effects.

Several potential kinases have been identified that target RLC_20_, including ROCK^12^, the ROCK-related citron kinase^13^, the zipper interacting protein kinase (ZIPK)^14,15^, integrin-linked kinase (ILK)^16,17^, the Inhibitor κB kinase 2 (IKK2)^18^ and the PIM3 (Proviral Integration site for Moloney murine leukemia virus) kinase.^19^. These kinases have been shown to phosphorylate both Ser19 and the adjacent Thr18 *in vitro*. However, the specific physiological implications of *in vitro* studies are not fully understood. There is evidence that PIM/ZIPK contribute to contraction and maintenance of vascular tone, and that inhibitors of these kinases can reduce BP in mouse models.^19^ Further, IKK2 deficient mice have been found to have decreased aortic contractile responses and reduced responses to hypertensive vasoconstrictors, suggesting a potential role in regulating vascular function and BP.^18^

The p90 ribosomal S6 kinase isoform 2 (RSK2), encoded by the gene *RPS6K3*, was another kinase observed to phosphorylate MLCK *in vitro* on Ser19.^20^ We recently demonstrated that RSK2 contributes to basal vascular tone, myogenic vasoconstriction of resistance arteries and blood pressure (BP).^21,22^ Our research revealed that this is mediated by direct phosphorylation of RLC_20_ as well as phosphorylation of the Na^+^/H^+^ exchanger, NHE1, which leads to an alkalinization of the SM cytosol and increased cytosolic [Ca^2+^], activating MLCK and/or other Ca^2+^-dependent kinase(s).^22^ RSK kinases (four isoforms) are atypical in that they have two catalytic domains: the functionally important N-terminal catalytic domain (NTKD) and the regulatory C-terminal (CTKD). The activation of RSK2 in cells is triggered by the phosphorylation of CTKD by ERK1/2 kinases on several residues, including Ser386, which creates a docking site for 3-phosphoinositide-dependent protein kinase 1 (PDK1). The latter phosphorylates and activates NTKD on Ser227.^23-25^ We observed that increased intraluminal pressure in resistance vessels to induce myogenic vasoconstriction results in phosphorylation of Ser227 of endogenous RSK2; moreover, vasoconstriction and Ser227 phosphorylation as well as RLC_20_ phosphorylation are blocked by RSK inhibitors BI-D1870 and LJH685.^21,22^ RSK2 signaling accounts for approximately 25% of the maximal myogenic constriction in normal mouse resistance arteries, implicating RSK2 as a significant player in vasoconstriction under physiological conditions. Accordingly, BP was significantly lower and the myogenic response suppressed in mice with global RSK2 knockout.^22^ Clearly, RSK2 may account for the contractility observed in the absence of MLCK, and could be a viable drug target for hypertension.

To understand the role of RSK2 in SM function, in a rigorous manner, it was necessary to conduct experiments utilizing a mouse model with complete *mylk1* knockout. Previously, we reported generation of a global *mylk1* knockout in mice, by disrupting promotor sequences located within intron 27.^26^ Although the embryos developed to full size, they died within 1-5 hrs after birth. The expression of all three *mylk1* products in these mice was abolished, as evidenced by western blot and polymerase chain reaction analysis. Unfortunately, at the time, we did not pursue further experimental characterization of the SM tissues from these embryos; we completed the work now and are reporting the full results. Further, we gained insights into the mechanisms responsible for contractions and RLC_20_ phosphorylation that occur in the absence of MLCK, including importantly the role of RSK2. Our data corroborate that RSK2 mediates these contractions. Additionally, we present compelling evidence that the apparent Ca^2+^-dependence of RSK2 is mediated by the upstream activity of the Ca^2+^ activated tyrosine kinase, Pyk2, which leads to the activatory phosphorylation of PDK1 and/or ERK1/2, with consequent upregulation of RSK2. This is consistent with previous findings about the physiological role of Pyk2 in SM.^27,28^ We also report that similar to MLCK, RSK2 binds to actin and to MLCK forming a signaling hub with PDK1 and Erk1/2 on the actin filament, ideally positioned to interact with the myosin RLC_20_.

Our findings provide further support of an important role for RSK2 signaling in vascular SM contractility, and provide rationale for the observed contractility of SM in tissues with *mylk1* knockout (*mylk1*^*-/-*^).

## Methods

### *Mylk1*^*-/-*^ mice

Briefly, a targeted construct, that includes a loxP flanked neomycin selection cassette was inserted in intron 27 of the *mylk1* gene; the construct was designed to allow Cre recombinase disruption of the telokin promoter sequence.^26^ This caused disruption of the entire gene resulting in a complete loss of MLCK (both 130- and 220-kDa isoforms) in homozygous animals. To make the most of the limited number of *mylk1*^*-/-*^ embryos, and the very small size of individual tissue samples, we harvested as many samples as possible from each embryo. This accounts for different SM tissues being used for the different assays. All animal studies were performed under protocols that comply with the Guide for the Care and Use of Laboratory Animals and were approved by the Animal Care and Use Committee at our institution.

### Genotyping

PCR analysis was performed on weaned pup’s and embryo tails or limbs using the following primers: Primers for MLCK: Telokin P1/telokin P3. P1 5’-AGA AGG AAA CTG AAG CCT GGA GAG GTC AAG-3’; P3 5’-ACT GTC AGC GTG TCC GAA GAT GTT CGG AGA ATG G-3’. Neo1/neo2: NEO1: 5’-CTT GGG TGG AGA GGC TAT TC-3’. NEO2: 5’-AGG TGA GAT GAC AGG AGA TC-3’.

For RSK2: F: 5’ TCT CTC CTG TAT TTC CTT TCA GG 3’. R: 5’ CT GAC CAC CAG GAA ACC ACA 3’. Neo170/Neo684: NEO170: 5’ TGA ATG AAC TGC AGG ACG AG 3’. NEO684: 5’ GC AAC CGA TGA GCA CTA TAA 3’

### *Mylk1*^-/-^ and *RPS6K3*^-/-^ primary aortic cells

Cultured smooth muscle cells were prepared from abdominal aortas from *mylk1*^-/-^ and WT E18.5 mice and 2-month-old WT, *RPS6K3*^*-/-*^ mice, cleaned of adventitia, cut into 1 mm^2^ pieces, and left undisturbed for 10 days in culture medium. The absence of RSK2 and MLCK proteins was confirmed by gel electrophoresis and western blotting. Cells were identified as SM cells based on the presence of SM myosin heavy chain expression detected on western blots.

### Histology

After fixation in buffered formalin, alcohol dehydration and paraffin embedding, 4-5μm sections of paraffin-embedded tissues were processed and stained with Hematoxylin and Eosin and mounted with Dpx mounting media.

### RLC_20_ phosphorylation

*Mylk1*^-/-^ and WT cells were serum starved for 24 h, then stimulated with serum for 5 min in presence of either LPA (0.5 μM) for 2.5 min, or with U46619 (1μM) for 30 s in the presence and absence of LJH685 (10 μM), added 15 min before serum. Protein was precipitated using 10% trichloroacetic acid (TCA)/ acetone^29^ and solubilized in 2x Laemmli buffer and prepared for gel electrophoresis. *Mylk1*^-/-^ and WT embryonic bladders (E14.5-18.5) were stimulated with LPA (0.5μM) for 5 mins, flash frozen and freeze-substituted in 10 % TCA/acetone at −80° C.^30^ Frozen strips were then washed in acetone and dried, homogenized in sample buffer in a glass-glass, hand-operated homogenizer and phosphorylation was estimated by western blotting. For imaging, the membranes were blocked with the Intercept^®^ Odyssey blocking buffer (Li-Cor). The ratio of phospho-RLC_20_ to total RLC_20_ was determined by immunoblotting with primary antibodies. Primary antibodies were visualized using secondary antibodies conjugated to either Alexa Fluor 680 (Thermo Fisher) or IRDye^®^ 800 (Li-Cor) for Odyssey imaging.

### Pyk2 and PDK1 phosphorylation

Aortic SM cells were serum starved for 24 h, then co-incubated in presence/absence of selective Pyk2 inhibitor PF4618433 (10μM) for 15 min. Thereafter, cells were stimulated in presence or absence of AngII (1μM) for 1 min. Protein was precipitated using 10% TCA and acetone and solubilized in 200 μl of 2x Laemmli buffer with 4% SDS. Cell were scraped off the plates and samples were boiled at 95 °C for 10 min then agitated overnight at 4 °C and prepared for gel electrophoresis or stored at −80 °C.

### Smooth Muscle Force Measurements

Helical strips of abdominal aorta, umbilical arteries or bundles from bladder taken from E14.5-18.5 embryos were cut and mounted on a bubble chamber^30^ in HEPES Krebs solution for force measurements in response to agonists or high [K]. pCa-force relationships were obtained following permeabilization with α-toxin by increasing [Ca^2+^] from pCa 6.3 to pCa 5.5 with each force normalized to the maximal force developed by the same strip at pCa 5.5.^31^ Ca^2+^-sensitization through the RhoA/ROCK signaling pathway was assayed by addition of 10 μM GTPγ S to α-toxin (500 U/ml) permeabilized bladder SM strips in pCa 6.3 solution.^32^ Once force had reached a plateau, 5 and 10 μM Y27632 was added to inhibit ROCK and relaxation was measured. pCa solutions and α-toxin permeabilization are described elsewhere.^30^ RSK2 signaling was determined by cumulative additions of carbachol in the presence or absence of 1 μM BiD1870 or DMSO diluent. Ca^2+^-independent force development in *mylk1*^-/-^ embryonic arteries and bladder was induced by the MLCP inhibitor calyculin A 100 nM in the absence of Ca ^2+^ (pCa 9.0).

### Smooth Muscle Collagen Fiber Preparation and Isometric Force Measurement

Contractility measurements were carried out on cultured *mylk1*^-/-^ embryonic aorta SM cells incorporated into collagen gels as previously described.^33^ In brief, cultured cells at a density of 2 × 10^6^ cells/ml were suspended in neutralized collagen in solution containing 1.2 mg/ml of collagen in PBS and transferred into a rectangular trough with a Teflon pole near each end. Culture medium was added after polymerization. After 14 days, the rod-shaped fibers were cut into strips and mounted for isometric force measurements on a bubble chamber. The fibers were stretched to 1.2 times slack length and equilibrated in the normal Krebs-bicarbonate solution, oxygenated and kept at 30 °C. Contractile responses to endothelin and the thromboxane analogue, U46619, were recorded.

### Cell Migration Assay

Proepicardia were isolated from E9.5 *mylk1*^-/-^ and WT littermates.^34^ To assay migration rates in the absence of MLCK, we used mesothelial cells (smooth muscle precursor cells) of the proepicardium, that undergo epithelial to mesenchymal transition and invade the myocardium to form coronary SMCs. The outgrowth of mesothelial cells, from proepicardia isolated from E9.5 *mylk1*^*-/-*^ and WT mice was recorded over 24 hrs. Edge speed was monitored over 24 hrs at 37° C.

### Immunoprecipitation Assays

Embryonic WT, *mylk1*^-/-^ and *RPS6K3*^-/-^ cells were serum starved for 24 hrs and stimulated with or without serum for 5 mins prior to centrifugation for 10 min at 4°C, 20,000 *g*. Cell lysates were prepared in 1% Triton X-100 TBS buffer. The supernatant was applied to Protein G Sepharose 4 Fast Flow beads (GE Healthcare) preincubated with or without MLCK mouse monoclonal antibody (Sigma Aldrich, St. Louis, MO, USA) and incubated at 4°C for 1 hour. After centrifugation, beads were washed (four times) with Tris buffer containing 0.1% triton X100 and proteinase inhibitor cocktail, Thermo, Invitrogen and solubilized in 2x Laemmli buffer and boiled at 95°C for 5 min. Extracted proteins were separated by SDS-PAGE at 70 V using 1.5 mm thick, 8% polyacryamide gel for western blotting. RSK2 phosphorylated at Ser227, PDK1 phosphorylated at Ser241, anti-ERK1/2 phospho-44/42 were detected using specific antibodies for these proteins. Primary antibodies were visualized using secondary antibodies conjugated to either Alexa Fluor^®^ 680 (Thermo Fisher) or IRDye^®^ 800 (Li-Cor) for Odyssey imaging.

### Proximity ligation assay (PLA Assay)

WT cells were grown in 10 mm confocal dish up to 70-80 % confluence (Corning), and serum starved for 24 h and then serum stimulated for 5 min followed by fixation, permeabilization and blocking. Thereafter, Duolink PLA assay kit (Sigma Aldrich) protocol, was followed for assessment of localization and co-association of RSK2 and MLCK. Two primary antibodies to RSK2 (anti-mouse) and MLCK (anti-rabbit) were used. PLA detects proteins within < 40 nm.

### Western blots, antibodies and reagents

Proteins were transferred to polyvinylidene difluoride (PDVF) or nitrocellulose membranes and blocked with Odyssey blocking buffer or 5%BSA, probed with primary followed by secondary antibodies in blocking buffer, and detected and quantified on the Odyssey system (LI-COR). The following antibodies were used: mouse monoclonal and rabbit polyclonal anti-actin (1:5000 WB; Sigma-Aldrich), MLCK (1:1000 WB; 1:250 PLA; 1:200 IP; Sigma-Aldrich and 1:500 IP; Abcam), goat polyclonal anti-RSK2 phospho-Ser227 and mouse monoclonal anti-RSK2 (1:500 WB; Santa Cruz, Biotechnology, Inc). Mouse monoclonal anti-RLC_20_ (1:1000 WB; Sigma-Aldrich. Rabbit polyclonal anti-RLC_20_ phospho-Ser_19_ (1:200 WB), anti-ERK1/2 phospho-44/42 (1:500 WB), anti-PDK1-phospho-Ser241 (1:500 WB), anti-Pyk2 phospho-Tyr402 (1:500) from Cell Signaling. For triple westerns proteins were separated on 1.5 mm thick, 10% polyacryamide gel at 70 V for 3 h, transferred on 0.2 μm PVDF membrane at 100 V for 2 h in transfer buffer containing 25 mmol l^−1^ Tris-HCl, 192 mmol l^−1^ glycine, 1% SDS and 20% methanol. Membrane was blocked in 5% BSA in 0.01% TBST, cut at 75 kDa molecular mass marker and incubated overnight with p-Pyk2 Tyr402, p-PDK1 Ser241 antibodies at 4°C. Signal was enhanced using a triple western blot protocol with biotin conjugated goat antirabbit IgG and Alexa Fluor 680 conjugated S-32358-streptavidin (1:100,000)^35^ and 31820-Goat anti-rabbit IgG (H+L) secondary antibody Biotin IgG (1:100,000) from Thermo Fisher Scientific were used.

### Statistical analysis

Data are expressed as mean ± SEM and analyzed by using either one-way ANOVA or two-way ANOVA followed by Bonferroni’s post hoc test. The *P*-value of less than 0.05 was considered significant. Graphs were plotted using Graph Pad Prism version 9 software (San Diego, California, USA).

## Results

### Generation of MLCK/telokin KO mouse

While generating the telokin-null mouse by homologous recombination^31^, we found that insertion of the full-length targeted construct into this locus, including the loxP flanked neomycin selection cassette, resulted in complete disruption of the *mylk1* gene expression in the absence of Cre recombination.^26^ The mice died shortly after birth. At E14.5 the litters (n=125) showed close to the expected Mendelian inheritance ratio of 1.5 -/-:4.8 +/-:1 +/+ with a normal average litter size of 8. Embryonic SM tissues or cultured aortic cells showed a complete absence of all products of *mylk1*, as evidenced by western blotting (Fig. 1A) or PCR (data not shown). This KO mouse will subsequently be referred to as *mylk1*^-/-^. Loss of telokin, a non-kinase SM-specific 17 kDa protein, identical to the C-terminus of MLCK is phosphorylated by PKA/PKG. ^31,36,37^ Phosphorylated telokin binds to the phosphorylated myosin phosphatase facilitating its binding to phosphorylated myosin and promoting myosin dephosphorylation.^38^ The lack of telokin in the *mylk1*^*-/-*^ embryos and SM cells is expected to have only a minor contribution to contraction events, if any, as cyclic nucleotide stimulation was not used and telokin expression is fivefold lower in aortae SM compared to Ileum.^31^

**Fig.1:**
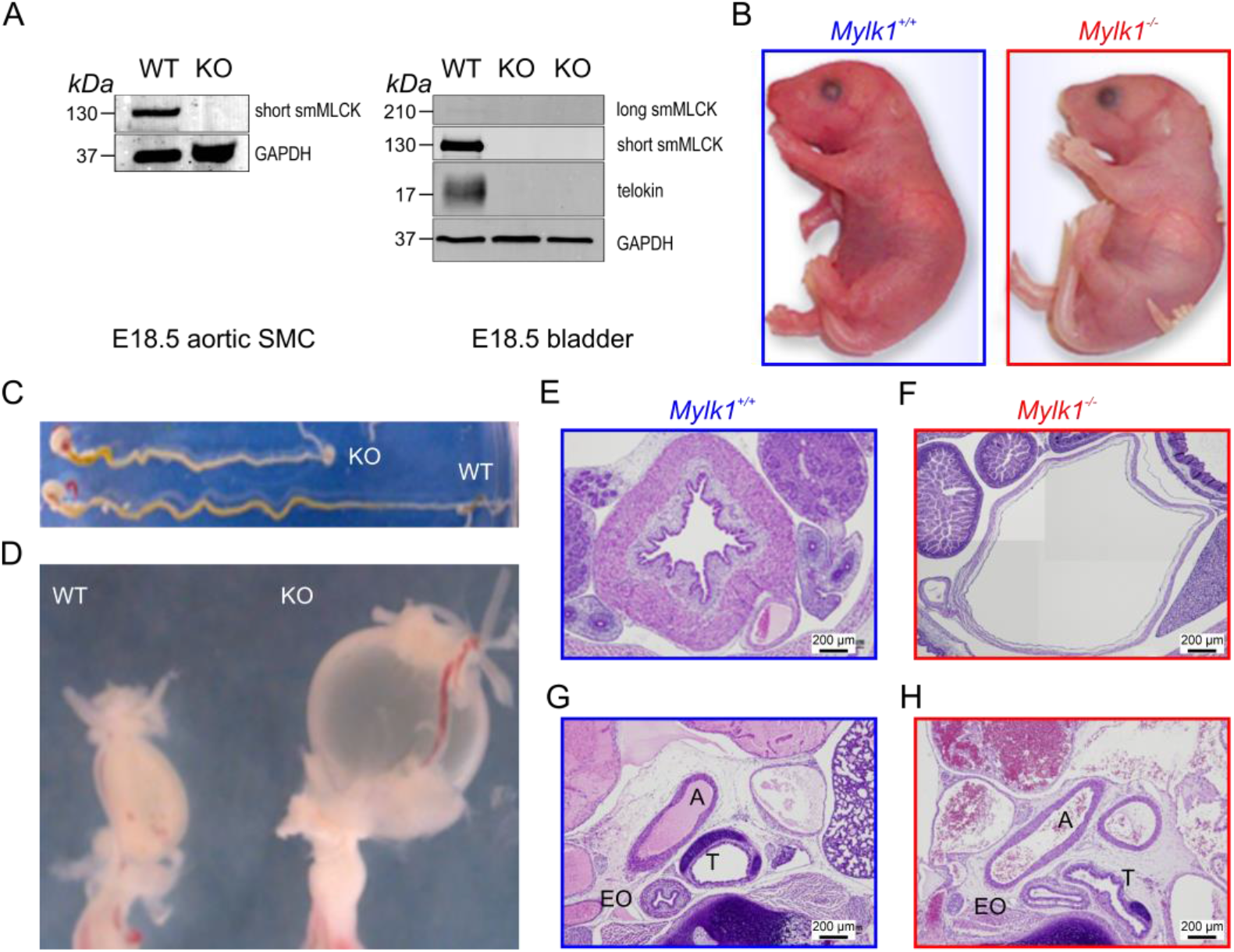
Long and short MLCK and telokin protein expression and phenotypic differences in *mylk1*^*-/-*^ and WT littermate mice. (A) Western blots showing complete loss of long and short MLCK and telokin protein in aortic and bladder SM from E18.5 *Mylk1*^*-/-*^ mice. Proteins were subjected to SDS-PAGE followed by western blotting with the anti – smMLCK (1:10000) and anti – telokin (1:1000) antibodies and fluorescent Alexa Fluor 680 anti – mouse secondary antibody (1:15000) and visualized by ODYSSEY Imager. GAPDH served as a loading control. (B) Postnatal pups. *Mylk1*^*-/-*^ died shortly after birth. *Mylk1*^*-/-*^ pups are typically paler, the length of the gut, stomach to rectum is shorter (C) and the bladder is markedly enlarged (D, E, F). Histology shows other dilated smooth muscle containing organs in transverse sections of E18.5 day embryos, stained with hematoxylin and eosin (G and H). A=aorta, T=trachea and EO = esophagus.

### Phenotypic changes in mylk1^-/-^ tissues

In general terms, the *mylk1*^-/-^ embryos were physically indistinguishable from WT and heterozygous littermates (Fig. 1B). Pups at birth, or embryos at E18.5 following dissection from the uterus, are pale in skin color as compared to both wildtype and heterozygous littermates. In addition, newborn KO animals tended to gasp for air within two hours following delivery and prior to death. However, the lungs do inflate and there is closure of the ductus arteriosus following birth determined by visualization under a light microscope (data not shown). Histological analysis showed dilation of bladders, aortae, trachea and esophagus (Fig. 1 D,E) consistent with a loss of SM tone in the absence of MLCK. Additionally, the length of the intestines from stomach to anus was shorter in the E18.5 *mylk1*^-/-^ embryos compared to wild type (Fig. 1B *p*<0.0005). The bladders were always engorged, reflecting the loss of basal SM tone and bladder sphincters were patent (Fig. 1D).

### Mylk1^-/-^ smooth muscle retains contractile activity

The initial phasic component of the contractile response to high [K^+^] was approximately 15 times greater and significantly faster in the WT than the *mylk1*^-/-^ embryonic bladder and umbilical arteries (Fig. 2A,B). This suggests that MLCK makes a significantly larger contribution to high [K^+^] -induced depolarization than the compensatory kinase(s), which exhibit slower kinetics. However, the compensatory kinase(s) in the absence of calcium is capable of inducing maximal forces equivalent to WT SM when exposed to the myosin phosphatase inhibitor, calyculin A, which allows for full RLC_20_ phosphorylation (Fig. 2A). When SM was permeabilized with α-toxin, sensitivity to increasing intracellular [Ca^2+^] was not different between WT and *mylk1*^-/-^, although the maximal force was lower in the *mylk1*^-/-^ (Fig. 2C). The small GTPase RhoA, known to significantly contribute to contraction of SM through activation of ROCK and inhibition of MLCP^3^, was also investigated to determine whether this signaling pathway is altered in m*ylk1*^-/-^ mice. Addition of the non-hydrolyzable GTPγS—to generate active GTPγS*RhoA—to α-toxin permeabilized MLCK^-/-^ and WT SM, in the presence of pCa 6.3, contracted the muscles and the contraction was reversed by subsequent addition of the ROCK inhibitor Y27632 (Fig. 2D). Furthermore, embryonic cultured *mylk1*^-/-^ aorta SM cells, when incorporated into collagen gels, contracted in response to agonists known to activate RhoA signaling as well as mechanisms that increase cytosolic [Ca^2+^] (Fig. 2E).

**Fig.2:**
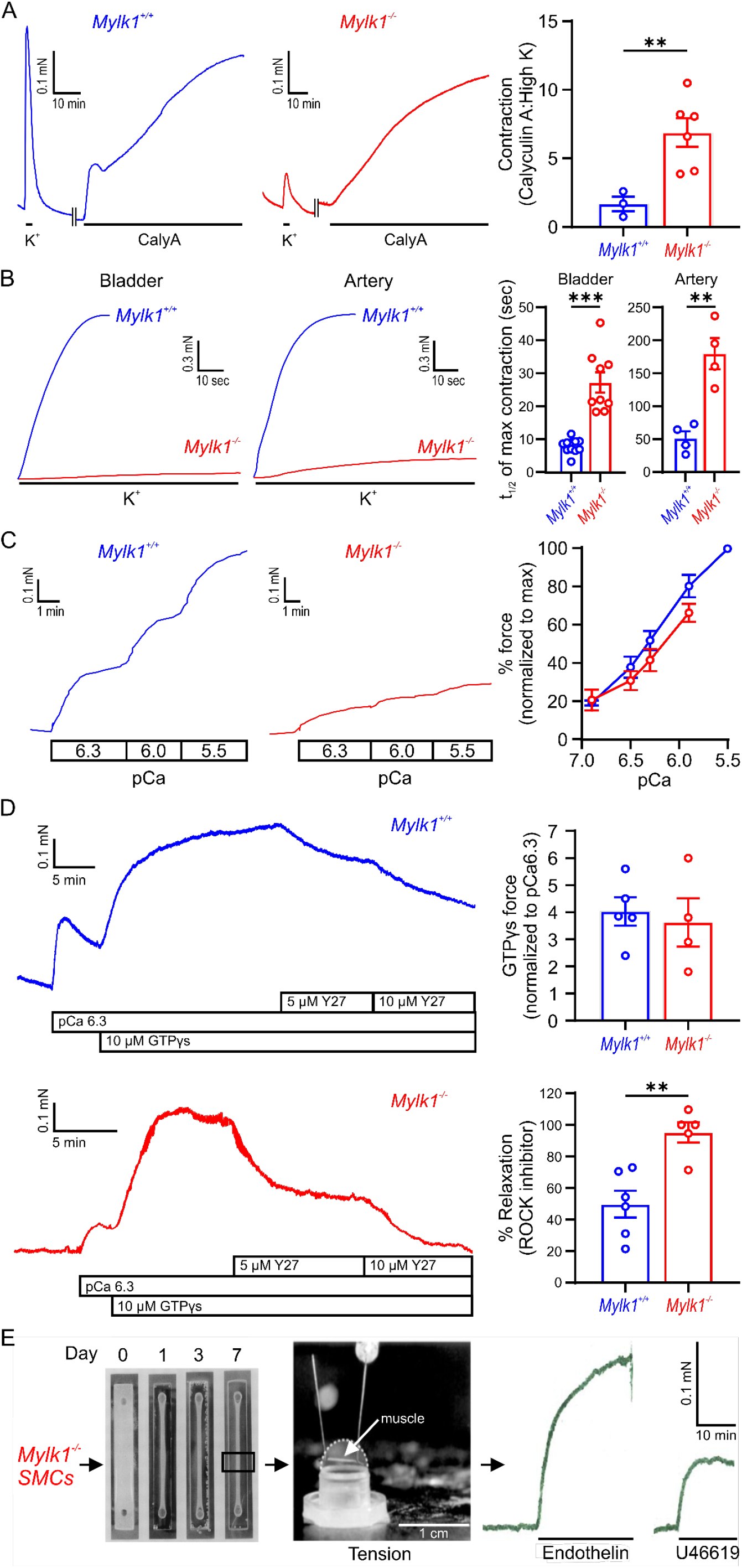
Assays demonstrating that *Mylk*^*-/-*^ SM retained contractile activity: (A) Umbilical arteries from WT and *mylk1*^*-/-*^ E18.5 embryos developed different magnitudes of force in response to high [K]. Following permeabilization with α-toxin, phosphatase inhibitor calyculin A (100nM) induced comparable magnitudes of maximal force in the *mylk1*^*-/-*^ and the WT littermate vessels in pCa>8 EGTA buffered solution. Ratios of calyculin A : high [K] are shown in the graph. (B) Different rates of contraction in response to high [K] occur in WT compared to *mylk1*^*-/-*^ in both bladder and umbilical arteries. Graph, the half-time of high [K] contractions in *mylk1*^*-/-*^SM strips was significantly greater than wildtype in both bladder (p<0.0001) and arteries (p<0.05). (C) *Mylk1*^*-/-*^ SM contracts in responses to increasing [Ca^2+^]_i_, in α-toxin permeabilized bladder E18.5 *mylk1*^*-/-*^ vs WT SM strips. Left panel, pCa-tension responses. Graph, pCa-tension curve normalized to force at pCa 5.5 (100%). The magnitude of contractile force was less but the sensitivity to Ca^2+^ was not significantly different in *mylk1*^*-/-*^ vs WT bladder SM (n=7 and 5 biological replicates respectively). (D) Activation of the RhoA/ROCK signaling pathway by addition of 10 μM GTPγS to α-toxin permeabilized WT and *mylk1*^*-/-*^ bladder in pCa 6.3 solution. The ROCK inhibitor, Y27632, 5 and 10μM relaxed the GTPγS-induced contractions. Graphs showed that the magnitude of GTPγS-induced contractions did not significantly differ between *mylk1*^*-/-*^ and WT SM (n=4 and 5 biological replicates respectively) but that the Y27632-induced relaxation of GTPγS-induced contractions was greater in the *mylk1*^*-/-*^ SM compared to WT (*p*<0.01, n=5 and 6 biological replicates respectively). (E) SM cells cultured from *mylk1*^*-/-*^ aortae of E18.5 embryos were mixed with collagen, cast into chambers and cultured for 7 days. The gels shrink and spindle shaped SMCs orient longitudinally. Small strips cut from the cell/collagen fiber were mounted on a bubble chamber. Contractile force developed in response to endothelin or the thromboxane analogue in the absence of MLCK.

### Migration of proepicardial cells were slowed in mylk1^-/-^ mice

The mesothelial cells of the proepicardium migrate over the surface of the heart and undergo epithelial to mesenchymal transformation accompanied by expression of smooth muscle specific markers. They migrate into the myocardium and invest on the endothelial network to eventually form the coronary vasculature. Migration assays showed that this process occurs in the absence of MLCK, however it is slowed by 50% (Fig. 3).

**Fig. 3:**
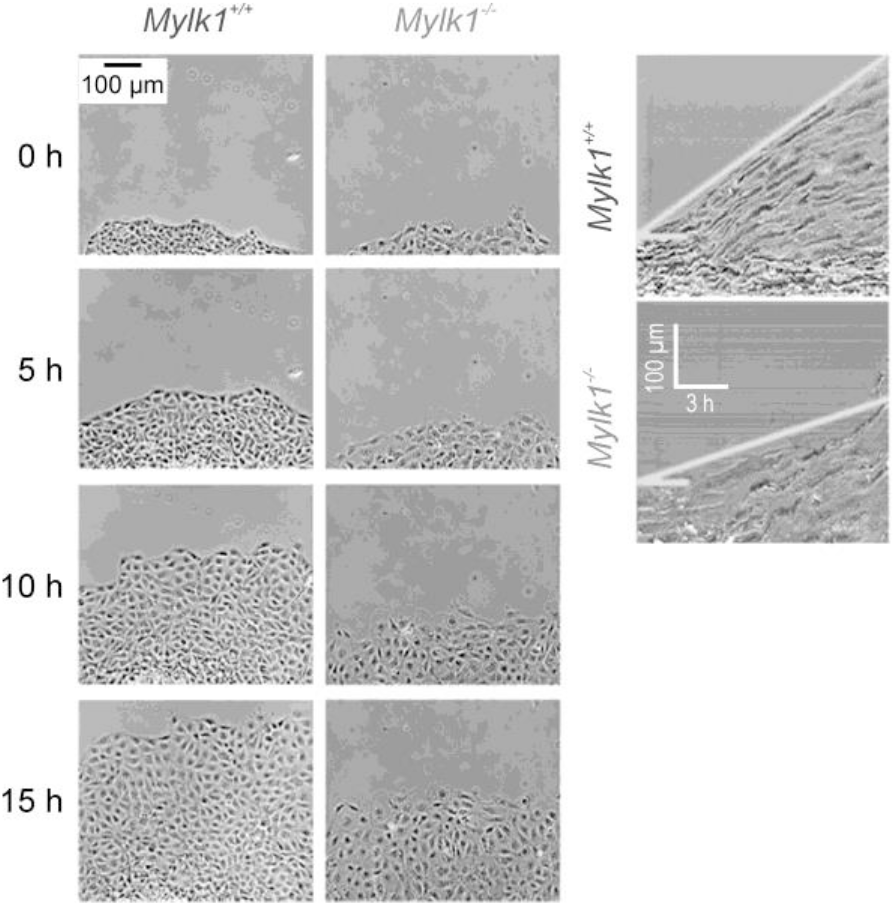
Migration of mesothelial cells from the proepicardium from *mylk1*^*-/-*^ mice were slowed. The outgrowth of mesothelial cells, (smooth muscle precursor cells) from proepicardia isolated from E9.5 *mylk1*^*-/-*^ and WT mice was recorded over 24 hrs. *Mylk1*^*-/-*^ cells migrated but the edge speed of migration was ∼50% slower in the absence of MLCK,) 0.19+/-0.06 vs 0.12+/- 0.04 μm/min, *p*<0.001, n=5 and n=4 biological replicates respectively.

### Myosin RLC_20_ phosphorylation occurs upon stimulation of mylk1^-/-^ SM and inhibition of RSK2 suppresses this RLC_20_ phosphorylation

Based on the ability of *mylk1*^-/-^ SM to contract, we next examined whether phosphorylation of myosin RLC_20_ —a necessary step for activation of myosin crossbridge cycling — occurs upon contraction of *mylk1*^-/-^ SM. LPA stimulation of cultured embryonic aortic *mylk1*^*-/-*^, *mylk1*^*+/-*^ *mylk1*^*+/+*^ cells resulted in an increase in RLC_20_ phosphorylation in both the presence and absence of MLCK (Fig. 4A). Stimulation of serum starved cultured aortic cells from *mylk1*^-/-^ SM with serum plus LPA or the thromboxane analogue, U46619 (Fig. 4B) significantly increased RLC_20_ phosphorylation, that was inhibited by RSK2 inhibitor, LJH685, a potent, ATP-competitive, and selective RSK2 inhibitor (Fig. 4B). We also established that our cultured *mylk1*^-/-^ cells were typical of SM cells as they expressed SM MHC, and actin and exhibited the ‘hills and valleys’ morphology typical of SM (Suppl Data Fig.1). As RLC_20_ phosphorylation is a key event for initiating crossbridge cycling in SM, the increased RLC_20_ phosphorylation in the *mylk1*^-/-^ SM demonstrates the presence of another kinase(s) capable of inducing RLC_20_ phosphorylation, accounting for the contractile capabilities of *mylk1*^-/-^ SM. Inhibition by the RSK2 inhibitor further supports our hypothesis that RSK2 is the kinase largely responsible for contractile function in the *mylk1*^-/-^ SM.

**Fig.4:**
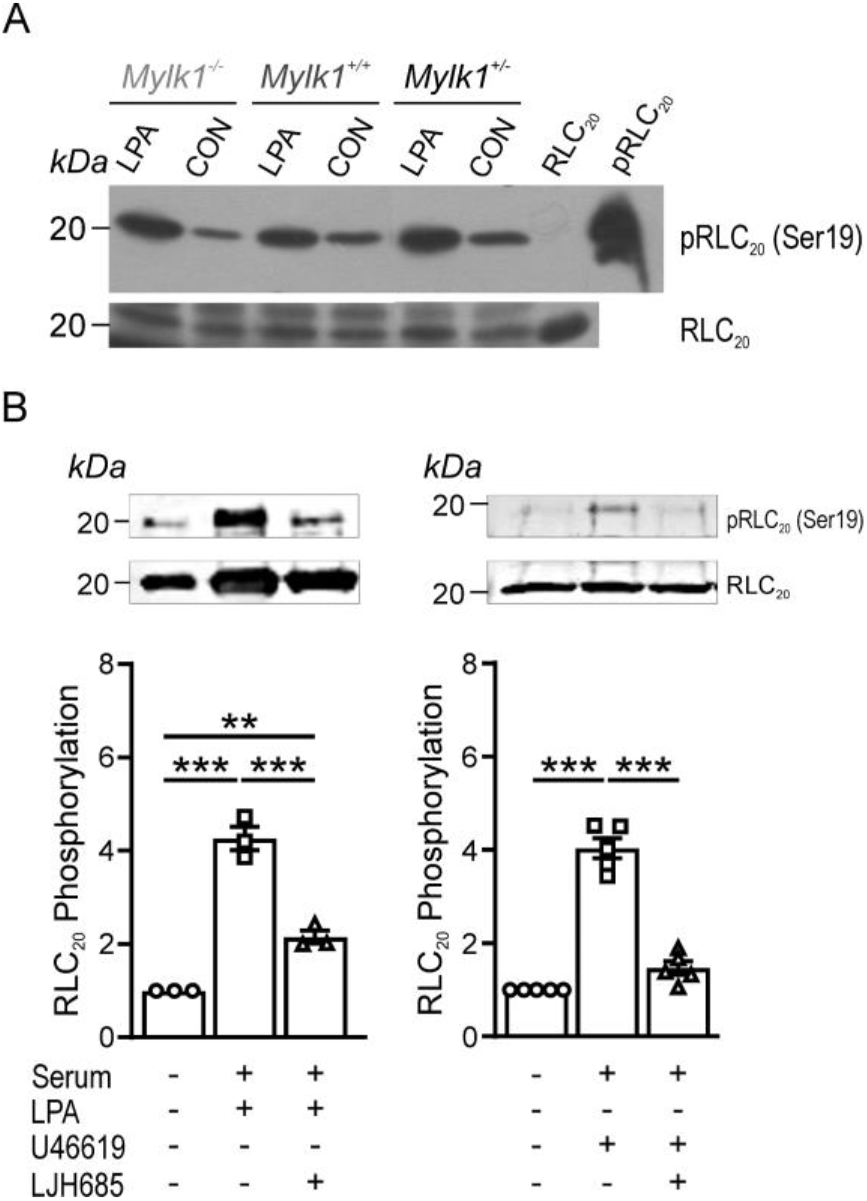
Myosin RLC_20_ was phosphorylated upon agonist stimulation in *mylk1*^-/-^ SM cells and was inhibited by RSK inhibitor LJH685. (A) LPA stimulation of cultured embryonic aortic *mylk1*^*-/-*^, *mylk1*^*+/-*^ *mylk1*^*+/+*^ cells resulted in an increase in RLC_20_ phosphorylation. Purified unphosphorylated and phosphorylated RLC_20_ protein served as controls. (B) RSK inhibitor, LJH685, significantly inhibited RLC_20_ phosphorylation in *mylk1*^*-/-*^ SM cells stimulated with LPA (*p*<0.001, n=3 biological replicates) or with thromboxane analogue, U47619 (*p*<0.001, n=5 biological replicates).

### Inhibition of RSK2 kinase suppresses carbachol-induced contractility of mylk1^-/-^ bladder SM

High [K] induced contractions in *mylk1*^-/-^ bladder SM were tonic, lacking an early phasic component observed in the WT muscles which preceded a tonic phase (Fig.5A, C). Contractile responses to increasing concentrations of carbachol were significantly inhibited in the presence of the RSK2 inhibitor BI-D1870 compared to the DMSO diluent in both *mylk1*^-/-^ and WT bladder SM, *p*< 0.0001 and *p*<0.003, respectively (Fig. 5B). Force was not completely abolished by BI-D1870 in the *mylk1*^-/-^ tissues.

**Fig. 5:**
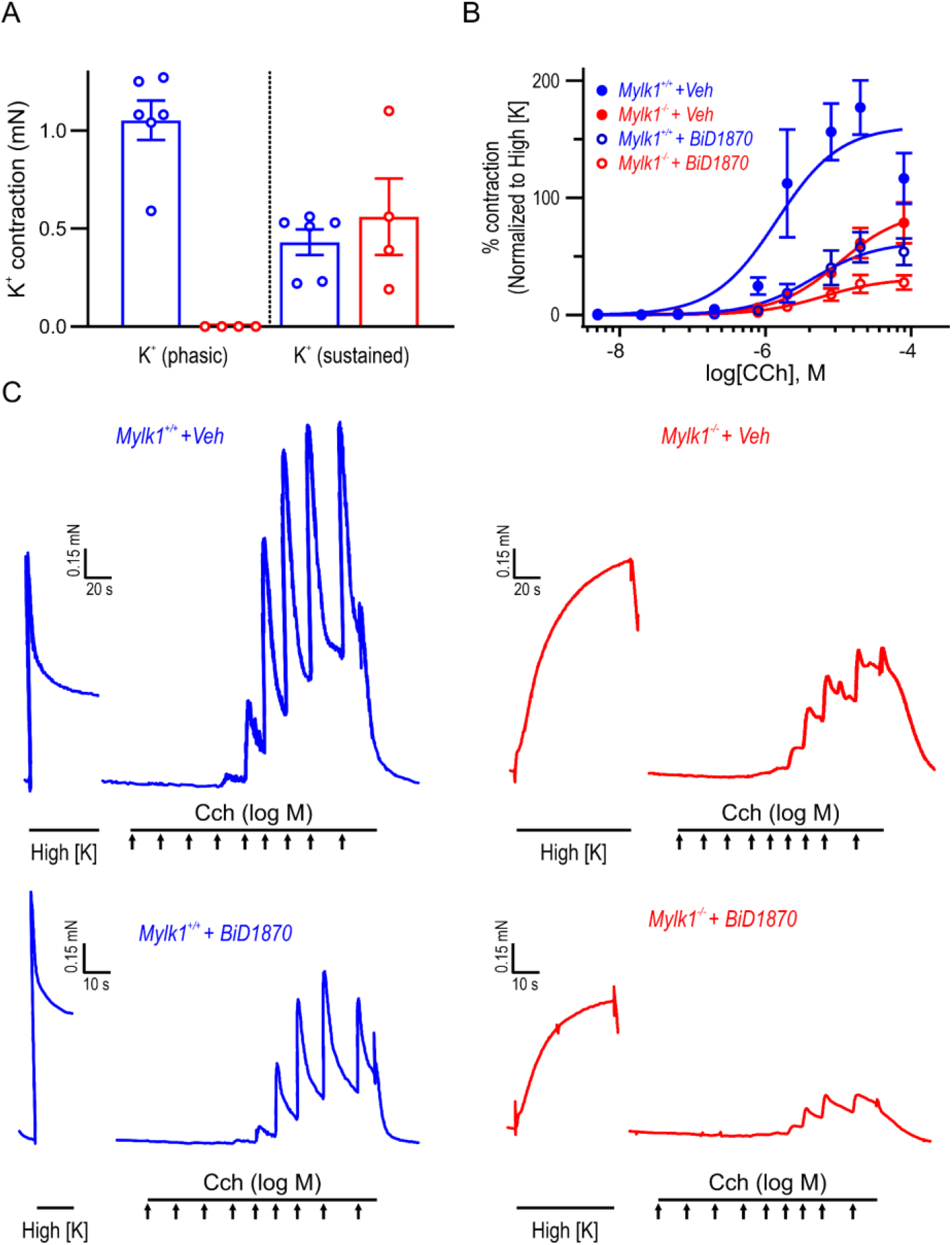
RSK inhibitor BiD1870 inhibited contractile responses to carbachol in *mylk1*^-/-^ and WT bladder SM. A) Summary of magnitude of high [K]-induced phasic and tonic components of contraction in WT (n=6 and *mylk1*^-/-^ (n=4) bladder SM. The high [K] phasic component was absent or small in the *mylk1*^-/-^ while the tonic components did not differ significantly in magnitude p=0.5. B) The contractile response to high [K] was recorded and following return to high [Na] Hepes buffered Krebs solution the muscles were treated with increasing concentrations of carbachol (Cch) in the presence or absence of the RSK inhibitor, BiD1870 (1μM) or equivalent volumes of diluent, DMSO. Traces shown in panel C. C) Contractile responses to increasing concentrations of Cch were normalized to the tonic high [K] contractions for each muscle and analyzed by 2-way Anova. WT DMSO vs WT BiD 1870: p<0.0001, *mylk1*^-/-^ DMSO vs *mylk1*^-/-^ BiD1870: *p*<0.003, n= 3 WT, n=2 *mylk1*^-/-^. Arrows=incremental Cch concentrations from 5×10^−9^ to 5 ×10^−5^.

### RSK2 colocalizes and co-immunoprecipitates with MLCK and actin upon stimulation in aortic SM cells

The protein ligation (PLA) assay indicates that RSK2 colocalized within less than 40 nm distance from MLCK and that this association occurred upon stimulation with serum and the thromboxane analogue, U46619 (Fig. 6A). RSK2^-/-^ and MLCK^-/-^ cells served as negative controls. This association of RSK2 and MLCK was confirmed in immunoprecipitation assays (Fig. 6B), where MLCK antibody pulled down RSK2, in WT but not *mylk1*^*-/-*^ SM cells. Beads alone served as a control and were negative (left panel) as was IgG (Supplementary data, Fig.2). MLCK immunoprecipitation also pulled down phosphorylated RSK2Ser^227^, RSK2, Erk1/2, phosphorylated PDK1Ser^241^, and actin (Fig. 2C). Serum stimulation significantly increased the association of MLCK with phosphorylated RSK2Ser^227^ with a tendency for an increase in phosphorylated Erk1/2, an upstream activator of RSK2.

**Fig. 6:**
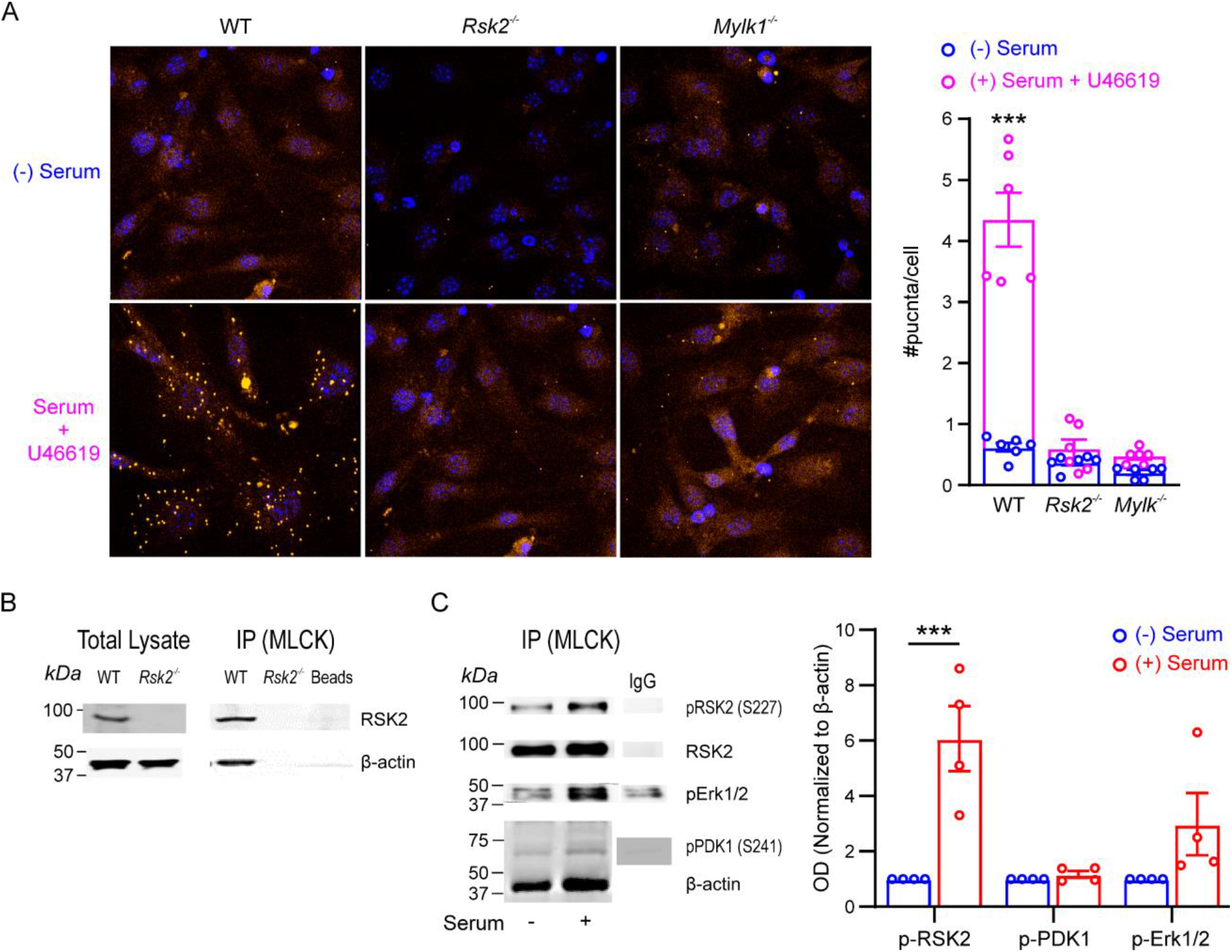
RSK2 and upstream activators, Erk1/2 and PDK1 associate with MLCK and actin in SM cells: (A) RSK2 and MLCK colocalized in the proximity ligation assay (PLA) in WT, but not RSK2^-/-^ and *mylk1*^-/-^ SM cells, in both serum starved or in the presence of serum plus the thromboxane analogue, U46619. Cells were labeled with rabbit monoclonal anti-MLCK antibody (1:200) and mouse monoclonal anti-RSK2 antibody (1:200), followed by secondary antibodies coupled to oligonucleotides (PLA probes). Following amplification of PLA probes in close proximity, complementary oligos coupled to fluorochromes hybridized to the amplicons and gave rise to discrete fluorescent puncta. The number of puncta/cell was significantly greater in the serum plus U46619 stimulated WT cells compared to non-stimulated (p<0.001 n=3 biological replicates. RSK^-/-^ and *mylk1*^-/-^ cells served as negative controls and had few puncta with or without stimulation. (B) MLCK immunoprecipitation (IP). RSK2 and actin were immunoprecipitated by MLCK in WT but not RSK2^-/-^ SM cells or beads alone. Neither RSK2, MLCK or actin were immunoprecipitated by IgG alone (Supplementary Fig.2 (C) representative western blot following MLCK IP from mouse WT aortic SM cells in the absence or presence of serum. RSK2, Erk1/2, PDK, phosphorylated-RSK2^S227^, phosphorylated Erk1/2 and phosphorylated-PDK1Ser^241^ as well as actin were immunoprecipitated by MLCK in both the presence and absence of serum (n=4 biological replicates). Serum conditions significantly increased the amount of phosphorylated-RSK2^S227^ immunoprecipitated by MLCK p<0.001, while there was an increased trend with phosphorylated-Erk1/2. Phosphorylated proteins were normalized to actin and 0 serum conditions taken as 1 for each phosphorylated protein.

### Ang II activates Ca^2+^-dependent tyrosine kinase, Pyk2 and the Pyk2 inhibitor (PF-4618433) inhibits activation of both Pyk2 and its downstream target PDK1

Agonist stimulation of aortic SM cells significantly increased Pyk2 phosphorylation, (Fig. 6), that was significantly inhibited by pretreatment with PF-4618433. Phosphorylation of the RSK2 downstream target, PDK1, known to phosphorylate S227 and activate RSK2, was also significantly inhibited by the Pyk2 inhibitor, as was phosphorylation of RSK2 S227.

## Discussion

Pathophysiology of vascular smooth muscle (VSM) constitutes a critical element of the etiology of hypertension (HT). Specifically, increased sustained vascular contraction, i.e. higher tone, leads to increased vascular resistance due to reduced vascular diameter and arterial remodeling, which is contingent on the ability of VSM cells to migrate.^39^ HT is a major risk factor for heart failure, myocardial infarction, stroke, vascular dementia, and other diseases, but the current pharmacotherapies—relying primarily on diuretics, β-blockers, Ca^2+^-channel blockers and renin-angiotensin pathway inhibitors—have limited success rates, leaving behind a significant patients refractory to treatments. It is not surprising that molecular mechanisms governing VSM contraction and cell migration are therefore actively studied in search of alternative drug targets. Most relevant cellular signaling pathways are regulated by protein phosphorylation, catalyzed by specific kinases. The latter are attractive drug targets, and many kinase drugs have been approved in recent two decades for clinical use, especially to treat cancers.^40,41^ and the field of design of kinase inhibitors for use in the clinic is highly advanced, with 68 approved by FDA as of 2022.^42^. Therefore, kinases involved in VSM regulation might potentially offer new avenues for the treatment of HT.

As described in the Introduction, recent studies showed that RLC_20_ is a substrate for phosphorylation by protein kinases other than MLCK1. The fact that complete global deletion of the *mylk1* gene does not prevent formation of embryos—even though they die after birth^26^ — provides compelling evidence that auxiliary kinases activating SM myosin are physiologically important in SM, and capable for compensating for lack of MLCK1. However, auxiliary kinases could not substitute for all MLCK functions as subepicardial coronary vessels from *mylk1*^*-/-*^ mice were irregular and migration of mesothelial cells from the proepicardium were slowed. This is most likely caused by the impaired migration of the fetal proepicardial cells that are precursors of coronary artery.^43^ Here, we expand the characterization of the *mylk1*^-/-^ global knockout, and demonstrate that SM tissues including bladder, fundus and blood vessels, as well as cultured SM cells, retain the ability to contract and migrate through the normal pathway, as evidenced by increased RLC_20_ phosphorylation in response to agonists (Fig.4). Importantly, addition of the MLCP inhibitor, calyculin A, to *mylk1*^-/-^ arteries and bladder in the absence of Ca^2+^ induces contractions of similar magnitude to those seen in normal arteries of similar size (Fig.2A). Therefore, under these conditions, the auxiliary RLC_20_ kinase is able to induce the full force potential of SM, and indicates that the contractile apparatus is intact. This kinase appears to be Ca^2+^-independent and operating in *mylk1*^-/-^ tissues, in the presence of intact RhoA/ROCK pathway (Scheme), which is also fully functional because embryonic SM tissues contract to GTPγS stimulation at constant [Ca^2+^] and inhibition of ROCK relaxes the GTPγS contractile responses (Fig.2D).

The percent relaxation due to the increased phosphatase activity, upon inhibition of ROCK, was significantly greater in the *mylk1*^-/-^ SM as the magnitude of contractile force depends on the ratio of kinase to phosphatase activity. RSK2 kinase activity is less and slower than MLCK, altering this ratio and allowing a preponderance of ROCK activated phosphatase activity. Likewise, the smaller and slower initial phasic high [K] contraction in the *mylk1*^-/-^ compared to the *mylk1*^+/+^ arteries reflects the depolarization induced rapid Ca^2+^ transient that activates Ca^2+^/CaM/MLCK present in the *mylk1*^+/+^ but lacking in the *mylk1*^-/-^ arteries (Fig. 2A, B).

The challenge now is to identify the most relevant protein kinase in VSM that might constitute a good target for therapy. We recently presented strong evidence supporting the notion that this role is played by the p90 ribosomal S6 kinase (RSK2), coded by the *RPS6KA3* gene, which has been hitherto recognized primarily for its role in brain development (*RPS6KA3* mutations cause the Coffin-Lowry developmental syndrome^44,45^) and cancer progression (it is a recognized target for cancer therapy, particularly the triple-negative breast cancer^46^). Specifically, we used SM tissues from global *RPS6KA3*^*-*/-^ mice, which are viable, to demonstrate the impact of deletion of RSK2 from SM. In the present, complementary study, we further explored the role of RSK2 in the SM tissues from *mylk1*^-/-^ embryos.

Arguably the most direct and compelling confirmation of our hypothesis in the present study is the fact that specific RSK2 inhibitors, Bi-D1870 and LJH685, inhibited carbachol-induced contractility in bladder of the *mylk1*^-/-^ embryos (Fig. 5), as well as RLC_20_ phosphorylation on Ser^19^ in *mylk1*^-/-^ SM cells stimulated with either LPA or U46619 (Fig. 4). Therefore, in the absence of MLCK1, contractility appears to be regulated primarily by the activities of RSK2 and the RhoA/Rock signaling pathways (Scheme, Fig. 7C). The low, residual contractility and RLC_20_ phosphorylation still observed in the presence of the RSK2 inhibitors may be due to incomplete RSK2 inhibition or to other kinases such as RSK1, integrin-linked kinase^16,47^,PIM/ZIPK^19^, or IKK2^18^. Nevertheless, our results support the notion that RSK2 is the long sought primary auxiliary contributor to SM contractility, independent of MLCK activity.

**Fig. 7:**
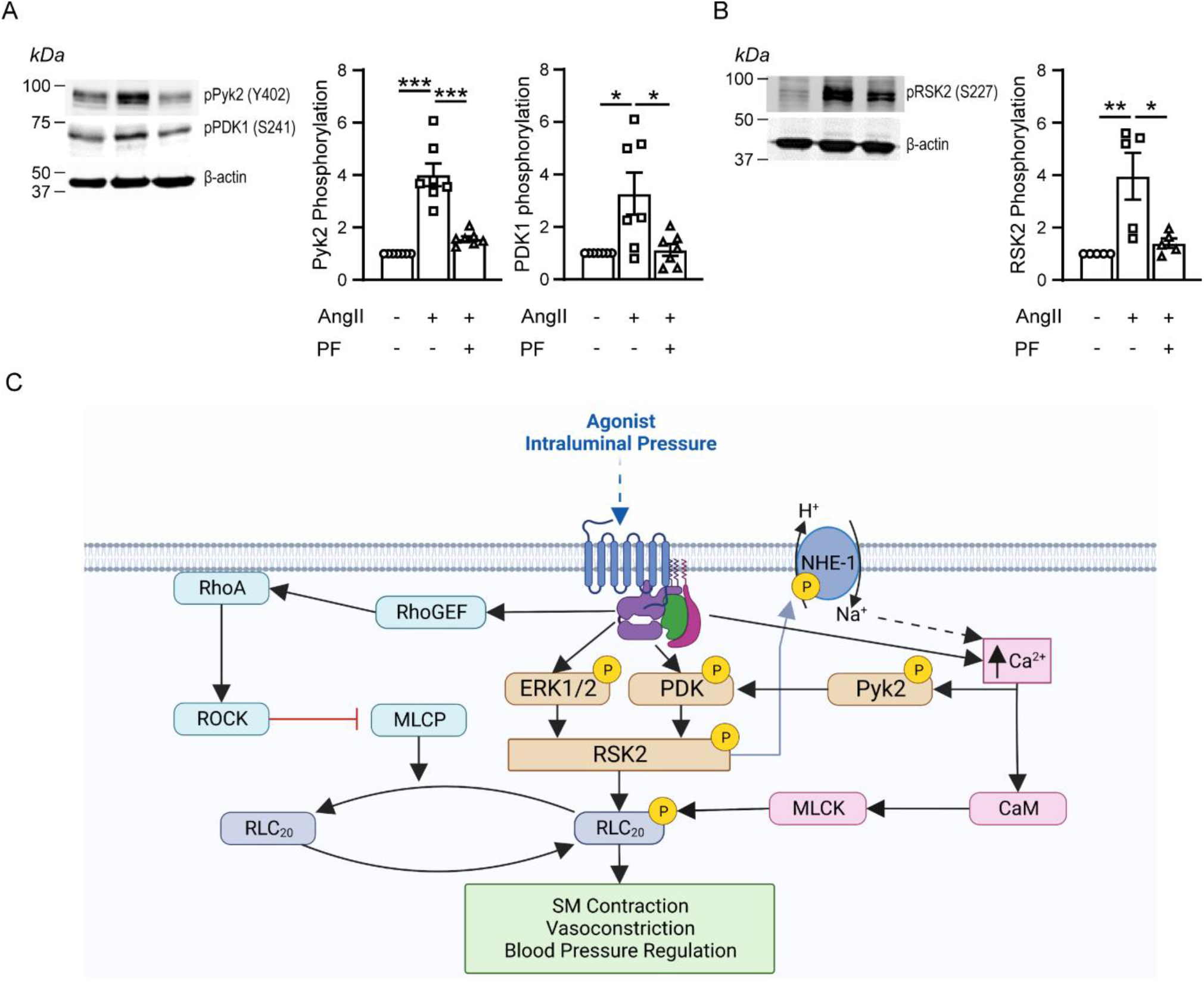
AngII stimulation increased phosphorylation of the Ca^2+^ dependent tyrosine kinase, Pyk2 and of its downstream targets PDK1 and RSK2 which was suppressed by Pyk2 inhibitor, PF-4618433. (A) Representative western blot and graphs showing significant AngII-induced phosphorylation of both Pyk2 (n=8 biological replicates, p<0.001) and PDK1 (n=7 biological replicates, p<0.01). The Pyk2 inhibitor, (PF-4618433), inhibits activation of both Pyk2 and its downstream target, PDK1 (n=8, p<0.001 and 7, p<0.01 respectively). (B) AngII increased RSK2 phosphorylation (n=5, p<0.05) that was significantly inhibited by PF-4618433 (n=5, p<0.01). Proteins normalized to actin. (C) Three signaling pathways regulating SM RLC_20_ phosphorylation, contractility, basal tone and regulation of blood pressure. Agonists, neurotransmitters or increases in arterial intraluminal pressure activate; 1) the canonical Ca^2+^ activated MLCK pathway. 2) the RhoA/ROCK pathway that phosphorylates and inhibits MLCP resulting in an increase in RLC_20_ phosphorylation. 3) the new RSK2 pathway that leads to activation of NHE-1 resulting in cytosolic alkalinization and increase in Ca^2+^ events. The increased Ca^2+^ feeds back to further activated the MLCK pathway. Activated RSK2 can also directly phosphorylate RLC_20_. Different stimuli may preferentially select for a given signaling pathway. Created with BioRender.com.

In normal SM cells, stimulation activates both MLCK and RSK2. In this case, as we have shown previously.^22^ RSK2 can both directly phosphorylate RLC_20_ and also activate the Na^+^/H^+^ exchanger NHE-1 resulting in an alkalinization of the cytosol, an increase in [Ca^2+^] and amplification of MLCK activity (Scheme, Fig. 7C). When these pathways dominate over MLCP activity, the net result is constriction, as in the case of myogenic vasoconstriction induced by agonist or intraluminal pressure, or in maintenance of basal tone in the resting state. It is however, of importance to note that not all three pathways are activated equally by different stimuli, or have the same time course.^48^ For example, thromboxane analogue U46619 is a specific potent activator of the RhoA signaling. On the other hand, depolarization with high [K^+^] is dominated by Ca^2+^/CaM/MLCK signaling as seen in Fig. 3A when comparing high [K^+^] responses in the arteries from the normal and *mylk1*^-/-^ mice. RSK2 signaling, in turn, contributes ∼25 % of the myogenic vasoconstriction response in resistance arteries, and contributes to resting blood pressure control.^22^ These multiple pathways provide both redundancy but also specialization depending on the stimuli, resulting in phasic or tonic type contractions.

Although RSK2 activity appears to be independent of MLCK, in normal SM cells MLCK immunoprecipitates with RSK2 and actin and increased with stimulation. The co-localization was confirmed in PLA assays, showing they are < 40 nm apart. MLCK has an actin-binding site in its N-terminus with its C-terminal kinase domain poised to reach across to interact with, and phosphorylate myosin heads to initiate myosin cross bridge cycling.^49^ We have previously reported that a fraction of RSK2 immunoprecipitates with actin filaments,^22^ while a soluble fraction is free to phosphorylate other substrates, such as the membrane associated Na^+^/H^+^ exchanger NHE-1. Interestingly, the two upstream activators of RSK2—PDK1 and ERK1/2— were also brought down with the immunoprecipitated complex of RSK2/MLCK and actin, suggesting a signaling hub on the actin filament poised to phosphorylate and activate neighboring myosin filaments. Few puncta were observed in the (-) serum PLA assay unlike the (-) serum immunoprecipitation assay where MLCK pulls down actin plus RSK2, Erk1/2 and PDK1. In this case, RSK2, possibly complexed to Erk1/2 and PDK1 may be bound to actin and not directly to MLCK. Thus, RSK2 may act independently of MLCK to phosphorylate myosin, as in the *mylk1*^-/-^ SM cells and/or associate with MLCK to modulate its activity. It remains to be determined, if the proximity of RSK2 to MLCK alters its activity and *vice versa*.

An intriguing question also remains as to how low (∼1 μM) concentrations of kinases tethered to actin filaments can phosphorylate ∼100 μM myosin heads. One reported possibility is that MLCK diffuses along actin filaments and enhances the rate of phosphorylation of SM myosin, as described by others.^50^ In that study, single molecules of fluorescent quantum dot labeled MLCK moved along actin filaments and preferentially located to areas in which myosin was not yet phosphorylated. The motion occurred only if the acto-myosin and MLCK-myosin interactions were weak. If the RSK2 associated with the MLCK/actin complex also moves with actin, this could similarly phosphorylate myosin heads if the RSK2 catalytic N-terminal domain can physically reach the myosin heads. Otherwise, the activated MLCK-associated RSK2 may regulated the activity of its MLCK partner.

One of our major concerns was that the results concerning [Ca^2+^] dependence of the auxiliary pathway were apparently contradictory. While the RSK2-mediated contractile force developed in *mylk1*^-/-^ SM in response to calyculin A was independent of Ca^2+^ (pCa>8), the permeabilized *mylk1*^-/-^ SM contracted in response to increased [Ca^2+^] (Fig. 2C). We previously reported that the pCa-force relationship in α-toxin permeabilized rabbit arteries was right-shifted and the maximal [Ca^2+^] response was inhibited by as much as ∼ 60% by either the RSK2 inhibitor BI-D1870 or the PDK1 inhibitor GSK2334470. These data suggest Ca^2+^-dependence of RSK2 and PDK1. AngII stimulation of SM cells from human resistance arteries has been shown to increase intracellular [Ca^2+^] and to induce a cytosolic alkalinization, which leads to increased activities of ERK(1/2) and several tyrosine kinases.^51^ These changes in [Ca^2+^] and alkalinization are consistent with AngII activation of RSK2 signaling^22^ and the tyrosine kinase activity suggest a role for another kinase. The non-receptor proline-rich tyrosine kinase, Pyk2 is Ca^2+^ dependent and phosphorylates PDK1 in AngII stimulated vascular SM ^28^. Pyk2 has also been shown to play a role in contraction of arterial SM. ^52,53^ Knockdown of PDK or Pyk2 inhibits LPA-induced phosphorylation of RSK2 and NHE3 activity in immortalized human colorectal adenocarcinoma cells.^54^ Using a highly selective inhibitor of Pyk2, PF4618433,^55^ we have found that AngII phosphorylates Pyk2 in aortic SM cells. PF4618433 inhibit both Pyk2 and its downstream target PDK1 as well as its downstream target RSK2 phosphorylation. Altogether, these findings are consistent with Pyk2 being upstream of PDK1 and RSK2 and accounting for the observed Ca^2+^-dependent component of RSK2 activity (scheme, Fig. 7C).

In conclusion, we present compelling evidence that RSK2 largely accounts for the contractile and phosphorylated RLC_20_ myosin observed in response to stimuli in SM from the *mylk1*^*-/-*^ embryos. These embryos have also allowed for the determination of the contributions of the RSK2 signaling to SM contractility in the complete absence of the canonical Ca^2+^/CaM/MLCK signaling. These findings support our previous studies showing the physiological importance of RSK2 signaling in the regulation of SM contractility and resting blood pressure. RSK2 contributes ∼ 25% of maximal force generated in response to increased intraluminal pressure in normal resistance arteries that regulate blood pressure and basal tone.^22^ All together, we propose that there are three major signaling pathways, driven by MLCK, RSK2 and RhoA/ROCK that regulate RLC_20_ phosphorylation and force in smooth muscle (Scheme, Fig. 7C). The magnitude of their contributions depends on the stimuli and the smooth muscle tissue.

## Acknowledgements

Supported by R01HL147555-01A1 to Somlyo A.V., Sonkusare S., and Z. Derewenda, R21NS118647 to Derewenda, Z., Somlyo, A.V. and S. Sonkusare and RO1HL147555-01A1S1 to R. Ayon. We thank Dr. Michael Walsh for advice on Pyk2 detection using triple westerns.

## Author Information

Jaspreet Kalra, University of Virginia

Mykhaylo V. Artamonov, Vaccine Research Center, NIAID, NIH, mykhaylo.

Hua Wang, Sentara Martha Jefferson Hospital

Aaron Franke, Brain Surgery Worldwide

Zaneta Markowska, University of Virginia

Li Jin, University of Virginia

Zygmunt S. Derewenda, University of Virginia

Ramon Ayon, University of Virginia

Avril Somlyo, University of Virginia

